# EcoKMER: One-stop shop for spatio-temporal metagenomic exploration using DataFed

**DOI:** 10.64898/2025.12.15.694490

**Authors:** August George, Damon Leach, Conner Phillips, Joshua Brown, Juliette M. Duncan, Blake Nedved, Owen P. Leiser, Noelani R. Boise, Ruonan Wu, Lindsey N. Anderson, Olga Kuchar, Margaret S. Cheung, David D. Pollock, Patrick Widener, Connah G. M. Johnson

**Author notes:** Co-corresponding authors: Connah G. Johnson, Patrick Widener. These authors contributed equally to this work (August George, Damon Leach, Conner Phillips).

## Abstract

Spatially distributed environmental sampling generates highly complex and multidimensional datasets illuminating key insights into microbial diversity, evolutionary-coevolutionary processes, and host-pathogen interactions. While these sampling methods generate high value datasets, dataset size, the dataset integration, visualization, analysis, and provenance tracking present significant bottlenecks to scientific discovery. To address this bottleneck, we developed EcoKMER, an R-Shiny front-end application designed to streamline metagenomic data accessibility and provide geospatial context to data in support of hypothesis-driven investigations into environmental sampling, supported by DataFed as its back-end data management platform. EcoKMER enables interactive visualization and filtering of harmonized metagenomic data and metadata using an interoperable approach, allowing users to extract spatially distributed sample-based metadata on top of environmental parameters such as geolocation, temperature, pH, for investigating ecological changes across time and space enhancing sample processing methods. As an example, we deployed this tool to track analysis of metagenomes from the organisms in the Salish Sea Estuary, consistent with existing community-accepted standards. Built on top of DataFed, a flexible and robust scientific data management system built for data lakehouse architectures, EcoKMER is positioned as a powerful tool to improve sampling strategy decision making, accelerate new insights for collaborative biological and environmental research, and fostering AI-ready analyses designed to enhance discovery and guidance for bioeconomic engineering. The published manuscript is available at https://www.osti.gov/biblio/3412938

## Background

Contemporary experimental workflows generate large quantities of data that are challenging to track and integrate with existing knowledgebases. Both large scale ecological sampling campaigns and automated laboratory workflows produce multiple rounds of data that require rapid analysis to guide practitioners towards tailored protocols and generate targeted hypotheses. This first triage analysis needs to provide quick and clear feedback to the experimenter that is integrated with contextual system metadata, such as equipment or protocol followed, to develop efficient targeted experiments. Facilitating such an analysis is a vital component of a quick turnaround modelling-experiment (ModEx) cycle used to improve data collection and insight generation workflows.

Both the rapidly developing field of laboratory automation and longitudinal environmental sampling campaigns produce a wealth of data that may be challenging to harmonize and visualize holistically [1]. These data sets will continue to grow, necessitating quick and clear human feedback in the loop operations. The structure, efficient serving, and provenance of these data will be vital for full adoption of agentic intelligence, which require effective labelling for dataset comparison and harmonization for the data to be “AI-ready” [2–4].

A key need for an analysis platform is to enable biological spatiotemporal visualization for the exploration of metadata with the support of a federated data management system. This platform would efficiently link environmental context for focused sampling efforts with a continuous feedback loop from collaborative researchers. Advancing our ability to analyze spatially distributed metagenomic samples is vital for understanding complex environmental processes, including host-pathogen interactions. However, the large-scale and multidimensional datasets generated during repeated environmental sampling efforts often present significant challenges in data processing, integration, and visualization. Without efficient tools and standards [5] to harmonize data and metadata, extract actionable insights, and enable hypothesis-driven investigations, valuable opportunities to reuse datasets for science discovery and experimental guidance may be overlooked.

## Findings

To address challenges in managing and visualizing spatially distributed metagenomic samples, we developed EcoKMER, an interactive R-Shiny application integrated to the DataFed management system [6]. We provide a schematic illustration of how EcoKMER is incorporated into a workflow in Figure 1. We deployed this tool to track the analysis of metagenomes from organisms in the Salish Sea Estuary. EcoKMER provides a streamlined solution for navigating diverse datasets generated from estuarine sampling efforts [7], aligning with FAIR (Findable, Accessible, Interoperable, and Reusable) principles while maintaining enhanced data provenance [8]. The user-friendly interface empowers researchers to filter and analyze datasets interactively based on spatiotemporal and environmental parameters such as geolocation, temperature, pH, and sample processing methodology. By enabling precise data selection, EcoKMER minimizes computational overhead by reusing data for analysis and triage, thereby improving the efficiency of data system investigations. The software integrates DataFed metadata management, data filtering, and exploratory analysis within a modular dashboard organized into two panels: Home and Data Analysis.

**Figure 1.**
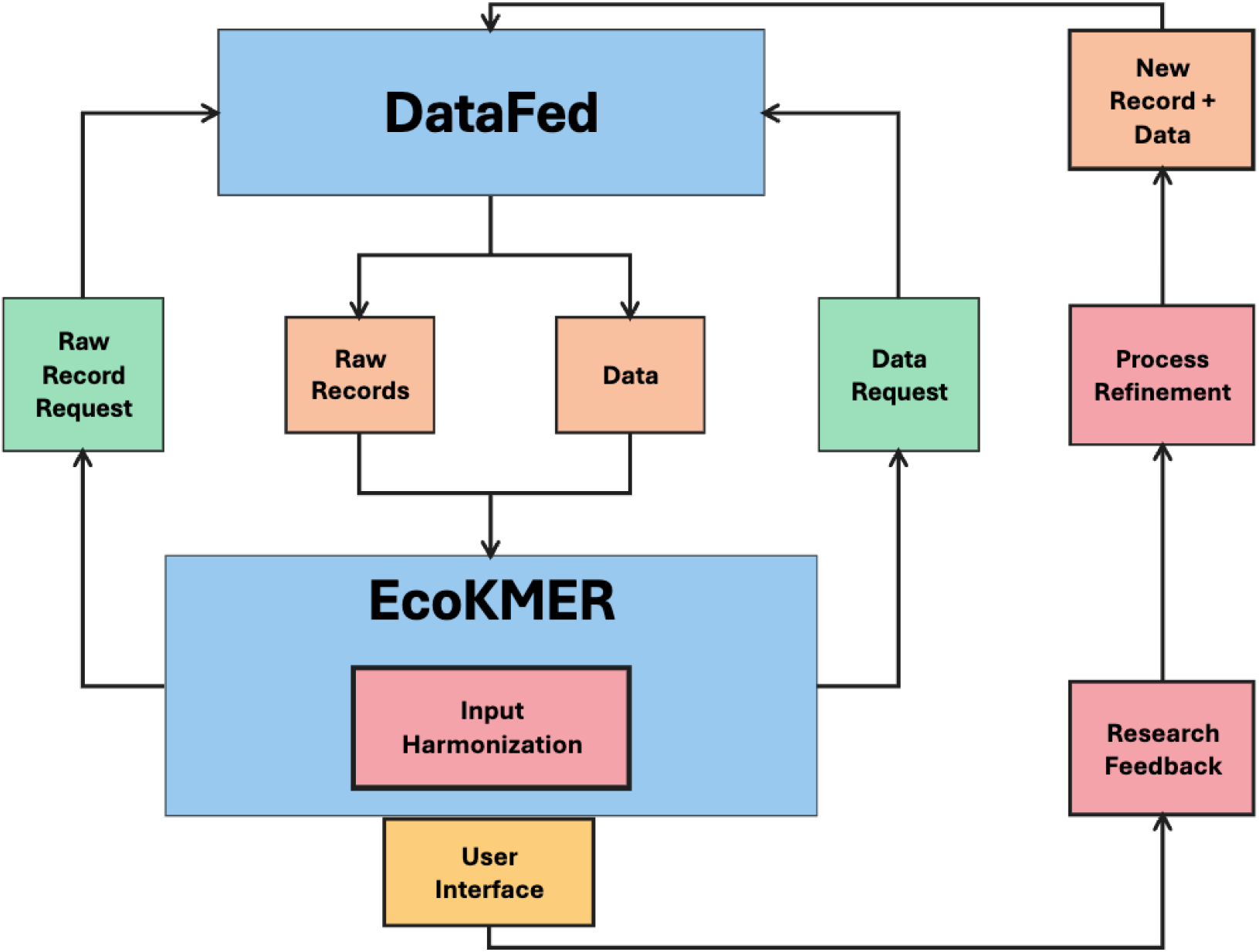
Demonstration of overall EcoKMER-DataFed data integration pipeline. New data records are created by a user using an appropriate metadata schema generated during the pipeline. These records are uploaded to DataFed and connected to related datasets for provenance tracking. These data records can be requested through the EcoKMER user interface where the metadata can be inspected and the full datasets can then be requested and served. Alternatively an offline-version supports the harmonization and triage of data without a DataFed connection.

EcoKMER provides a comprehensive framework to optimize workflows at the user interface level, separating data extraction and transformation processes from metadata exchange. The platform is ideal for rapid inspection of biological datasets for hypothesis generation and the ingestion of biological knowledge directly into system workflows, filling a critical gap in metadata exchange and visualization from large knowledge and resource databases. By streamlining processes and enabling flexible development while enforcing metadata schemas when uploaded to DataFed, EcoKMER enhances collaborative research efforts and improves accessibility to prepared datasets for focused investigations.

## EcoKMER Application Overview

EcoKMER’s interface (Figure 2) separates data ingestion from interactive analysis, enabling metadata-driven sample identification without downloading complete collections.

**Figure 2.**
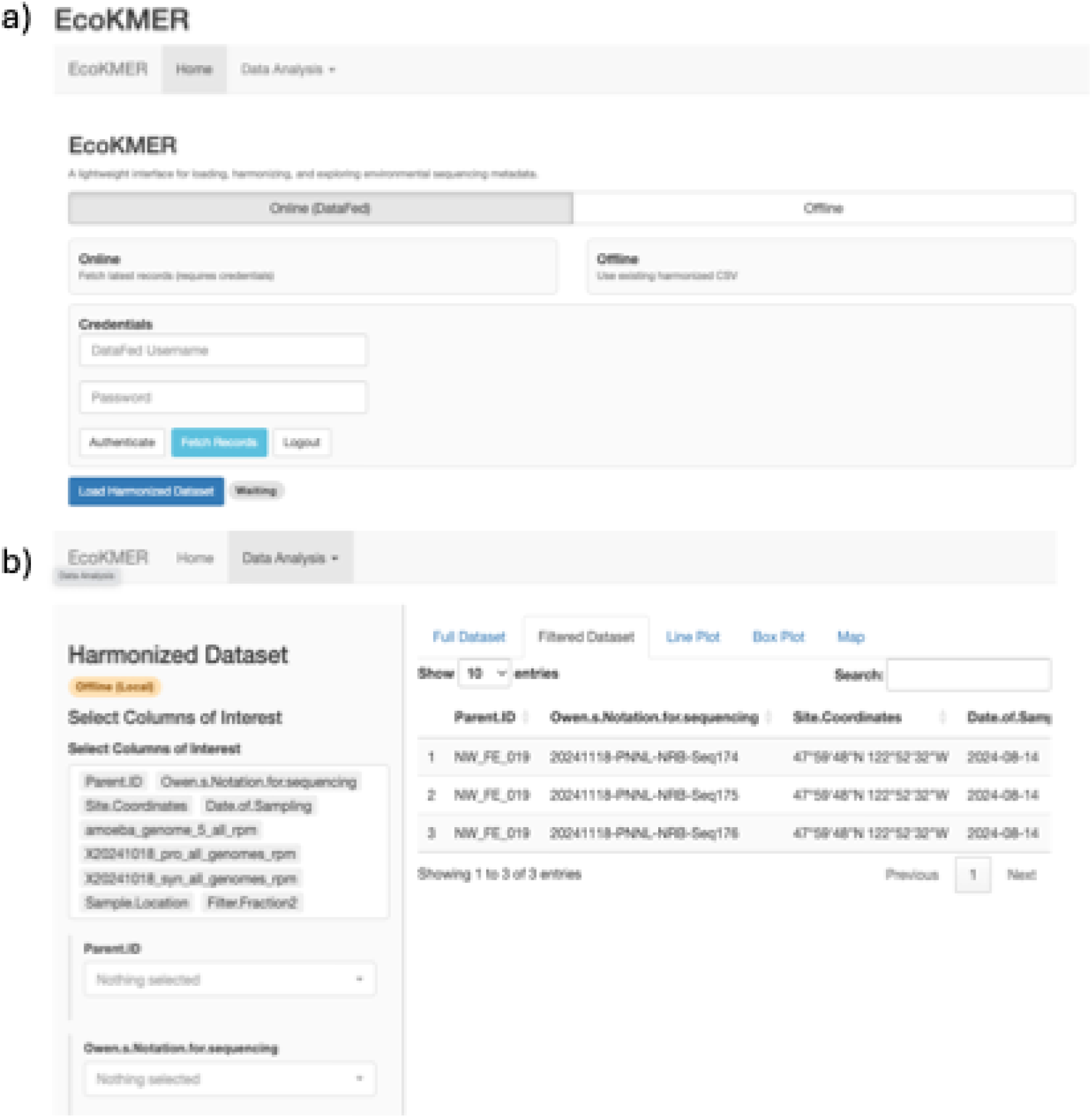
The EcoKMER application interface. Main components are (a) the home and (b) the data analysis panels. A user can log into the on-line DataFed repository on the home interface, to retrieve repository records, or select off-line mode, to ingest local data. Data from either source are analyzed and triage in the data analysis panel where harmonized data and plots can be formed.

### Home Panel

The home panel (Figure 2a) allows users dual-mode data access. On-line mode authenticates to DataFed [6, 9], retrieves project metadata, and merges collection-specific records into a harmonized dataset. After authentication, the application connects to the live DataFed infrastructure, providing access to the latest metadata and enabling the direct download of records through Globus [10, 11]. Offline mode loads locally stored metadata files, supporting instances where data cannot be transmitted freely or in-field analysis with limited internet connectivity.

### Data Analysis Panel

The data analysis panel (Figure 2b) uses a split-screen layout with filtering controls on the left with synchronized visualizations shown on the right. EcoKMER currently supports table views, box plots, line plots, and map visualizations. This approach leverages the complementary strengths of different visualizations: tables enable the verification of filter logic and identification of missing values, box plots show distributional differences across groups, line plots show trends between features, and maps contextualize spatial patterns. All the visualizations are interactive and dynamically updated based on the applied filters, supporting rapid data exploration without context switching.

### Spatio-temporal analysis

The map tab in the data analysis panel (Figure 3) displays georeferenced sample records with temporal color gradients. The markers are date-coded with dynamic visual clustering based on the zoom scale, preserving the median temporal signal through color averaging. This enables exploration of temporal patterns across scales, from regional trends to site-specific seasonal variation. EcoKMER transforms tabular sample metadata into interactive spatial visualizations without requiring GIS software or database expertise.

**Figure 3.**
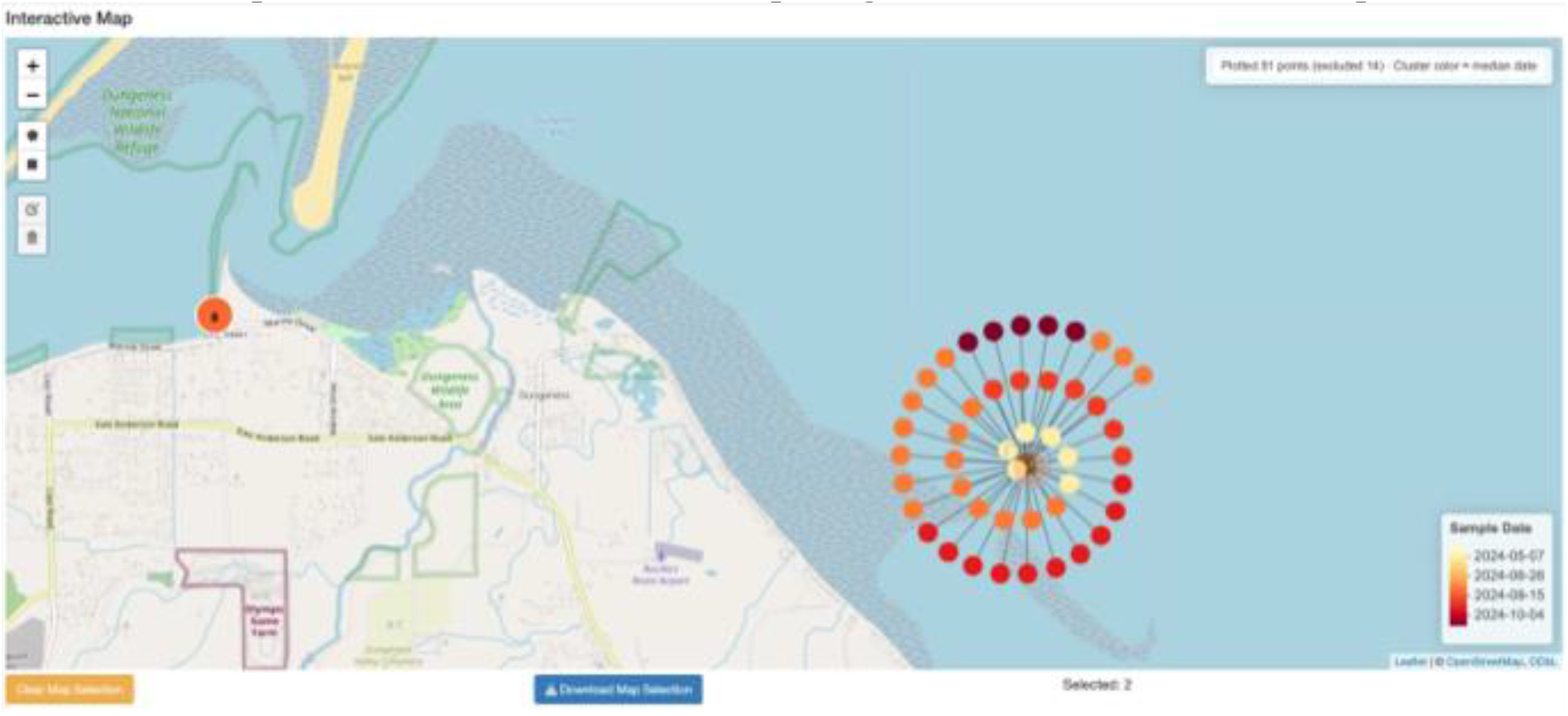
Map view along the example datasets across sampling events at site at PNNL-Sequim. This map demonstrates the spatial location of sampling events with dynamic visual clustering and time-based labels. This map provides spatial and temporal visualizations of the data records and used to provide contextual information regarding sampling events.

### Data selection and filtration

The filter section in the data analysis panel operates on the harmonized metadata records collected from the home panel. EcoKMER hierarchically operates on the metadata fields (i.e., columns) using interactive filters. Categorical variables use a dropdown selection, while numerical values use a range slider. Users can also select based on spatio-temporal features using the map view.

### Data retrieval and export

Users can export these selected subsets for further analysis. In offline mode, the filtered metadata is exported directly to a file. In online mode, users trigger a download of preprocessed sequence data from DataFed repositories based on their filtered metadata selection. This approach supports targeted data extraction that avoids unnecessary transfers from terabyte-scale collections common in metagenomic studies.

### Exemplar Analysis Visualization Workflow

The workflow is demonstrated by considering an exemplar sampling campaign for targeting cyanobacterial populations in the Salish Sea with amoeba as a large-celled organismal comparator [7].

## Federated Data integration with DataFed

EcoKMER integrates with DataFed and Globus to support collaborative metagenomic research across institutions in a platform of federated data. DataFed provides centralized metadata management while Globus coordinates data movement across distributed storage endpoints.

### Metadata validation and provenance

Metadata uploads to DataFed use JSON schemas defining column mappings, required fields, provenance relationships, and Globus transfer paths for associated sequence files. Schema validation occurs via external tool calls before upload to DataFed. In addition, DataFed supports dynamic server-side schema validation as data records evolve. DataFed’s typed provenance links connect sampling metadata to datasets. EcoKMER traverses these relationships to identify which artifacts correspond to the filtered metadata records. Graphical visualization (Figure 4) renders these provenance links as directed graphs, making processing workflows and data lineage immediately comprehensible.

**Figure 4.**
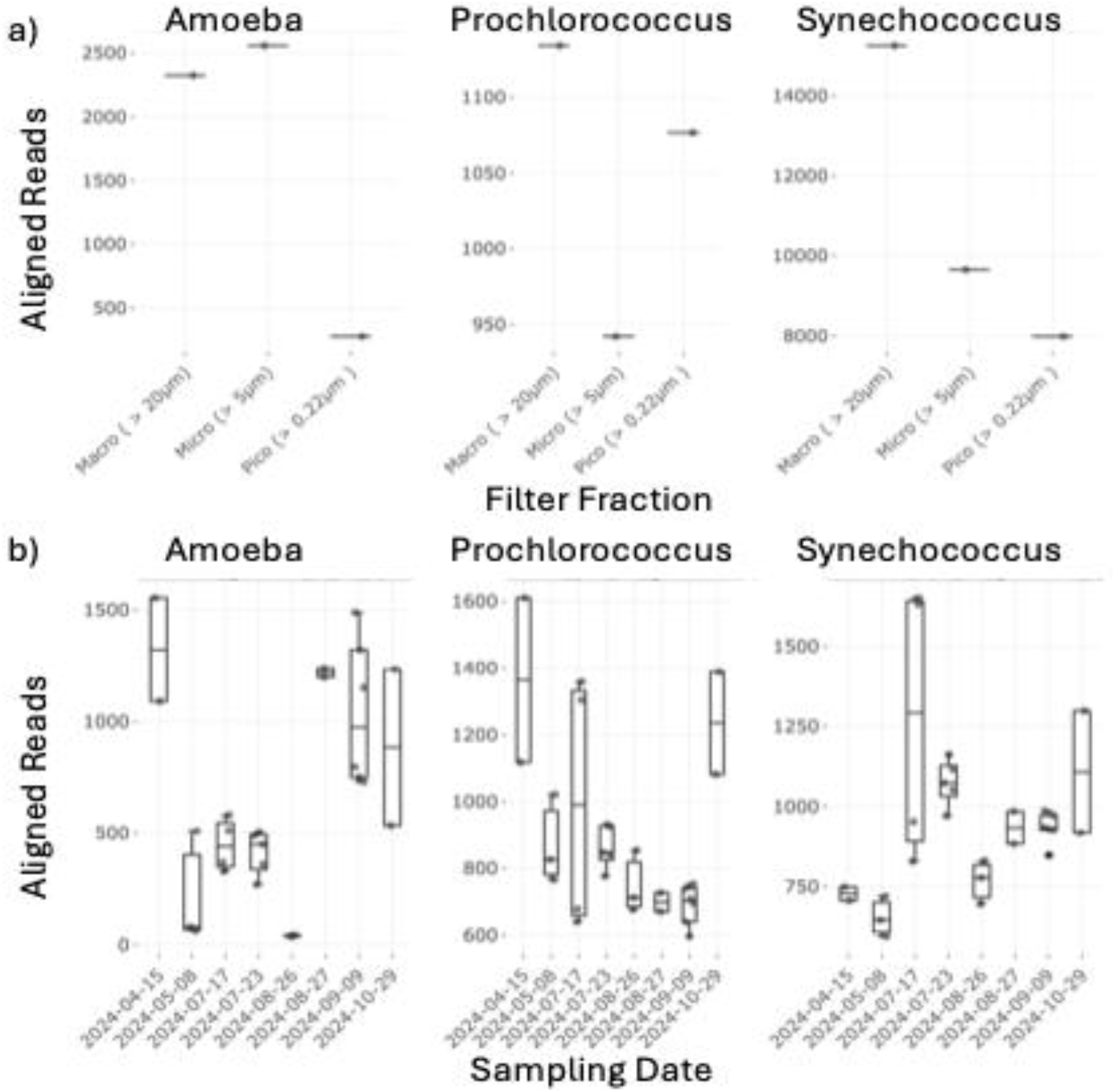
EcoKMER was used to analysis sampling events in the Salish Sea. The visualization capabilities of EcoKMER can be used for rapid analysis across the data records. The samples from a 6-month sampling campaign were filtered into different size fractions, sequenced, and aligned to provide metagenomic reads for the different size fractions (a) and sampling event times (b).

### Data storage and access

Metagenomic collections often span years and multiple sites, making full downloads impractical for exploratory subset analyses. EcoKMER integrates with DataFed’s federated architecture (Figure 5) to separate metadata filtering from file retrieval. DataFed repository administrators authorize collaborating users to access federated data on storage systems they managed. Authorized users can then query centralized metadata repositories, apply spatial-temporal filters interactively, and download matching raw files via Globus transfers to institutional endpoints (e.g. ORNL, PNNL, collaborator facilities). This metadata-driven approach significantly reduces bandwidth compared to downloading entire collections from distributed DataFed endpoints.

**Figure 5.**
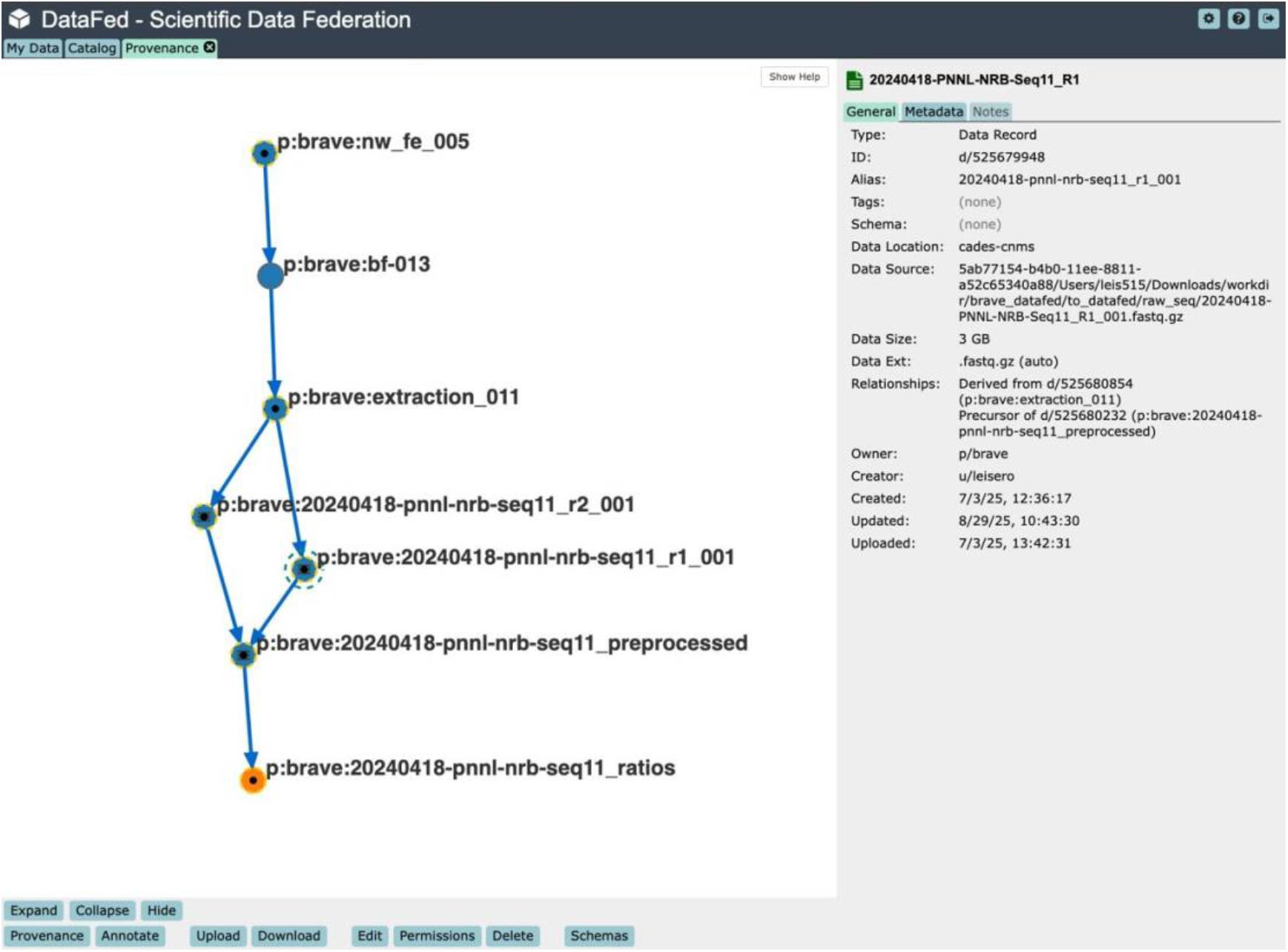
Demonstrating provenance and the record connections in DataFed. The on-line capability to leverage DataFed will ease this burden and allows user to track data provenance. DataFed backend supports interoperability and eases the burden on the user to need to supply their own data and make sure it conforms with EcoKMER’s input requirements, plus automatically ensures they have the most up-to-date data & data provenance tracking. DataFed provides both an interoperable backend communication protocol and a community information testbed for exchange of information.

**Figure 6.**
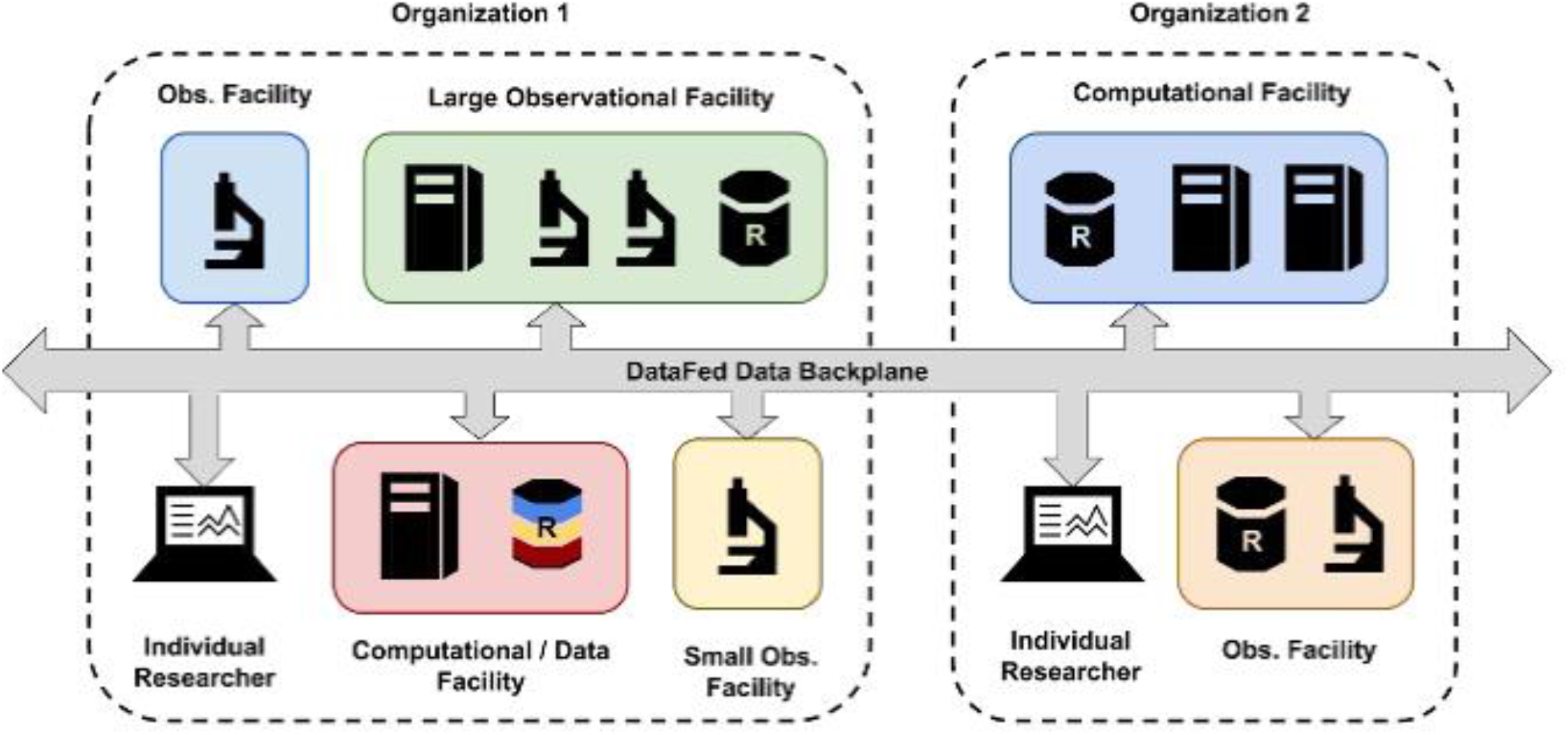
DataFed is data management software designed to provide uniform access to data spanning different projects. DataFed supports diverse collection/production modalities and cadences, and varying metadata needs.

## Software Architecture and Technical Design

This section describes the EcoKMER software architecture, data processing workflows leveraging DataFed for interoperability, and visualization components that enable reproducible exploratory analysis of metagenomic metadata.

### Distributed Data Architecture

The application bridges R-Shiny’s [12] reactive framework with DataFed’s Python API [6] via reticulate [13], enabling cross-language data operations. Python scripts implement distinct workflows: metadata retrieval from DataFed collections, file download orchestration via Globus endpoints, and dataset upload with dependency tracking. The application deploys as a local Shiny server.

### Data Harmonization

Heterogeneous input data undergo systematic transformation to standardize coordinates, dates, and numeric values for unified analysis. Coordinate parsing converts degrees-minutes-seconds (DMS) strings to decimal degrees. Date parsing uses sequential pattern matching to handle multiple formats. Missing and below-detection measurements are converted to a standard (i.e., R’s NA type) representation.

### Interactive Visualization

EcoKMER generates interactive maps via the leaflet package [14] and statistical plots using plotly [15], synchronized through Shiny’s reactive expressions. The statistical plots can export both vector and raster images.

### Reproducibility and Testing

Dependency management uses renv [16] for R packages, a requirements file for Python modules, and an instructive README for reproducibility. Quality assurance is provided by unit tests covering coordinate and date parsing, missing value handling, map and plot rendering, fixture schema integrity, and error handling.

## Conclusion

In this manuscript, we describe the development, functionality, and potential applications of EcoKMER, including exemplary use cases demonstrating its value for supporting scientific inquiry connecting field sampling to laboratory and computational analysis. EcoKMER complements existing metagenomic data platforms such as KBase [17] and NMDC [18] by addressing the data triage phase that precedes computationally intensive analyses. EcoKMER prioritizes rapid metadata exploration with selective data retrieval rather than comprehensive analysis pipelines. The included interactive spatial-temporal visualizations complement tabular metadata browsers typical of repositories like MG-RAST [19]. The dual-mode architecture supports both limited-connectivity field stations and multi-institutional projects.

By highlighting its data accessibility, visualization capabilities, and flexible analysis, we aim to illustrate EcoKMER’s transformative potential for integrative research. Enhanced by DataFed’s dynamic data management system, EcoKMER offers a flexible and extensible computational platform capable of integrating various data analysis tools to shorten the time of impact by allowing users to collaborate on-the-fly. Upcoming planned developments include plugins for sampling metagenomic analysis software such as NW-BRaVE’s Hydroplane [20], exploring plugins of other project codes, and integration of statistical testing capabilities to enable evolutionary analyses directly within the EcoKMER interface. Future enhancements will enable automated end-to-end workflow information exchange and seamless on demand search and discovery capabilities across data information resources supported by semantic metadata harmonization. Continued semantic refinements will support structured information exchange of experimentally informed computed workflow metadata defined by domain expert researchers. Continuous metadata schema extensions and refinements encourage version-controlled data community exploration and reuse of diverse project data types (e.g., simulated workflow executions, etc.) for driving new insights about host-pathogen interactions while maintaining model schema interoperability with standardized BER cyberinfrastructure frameworks such as NMDC. Combined with enhanced extraction of contextual information using large language models for metadata standardization and interpretation of annotations [21], these enhancements will advance the EcoKMER platform toward its goal of becoming a comprehensive, user-friendly “one-stop shop” application for data triage, remaining compatible with future opens science community suggested standards and fostering collaborative research that shortens the time of impact from field to laboratory.

## Data availability

All code and example input and output data are available at github.com/NWBRaVE/EcoKMER

Data presented in this work can be accessed on DataFed, https://datafed.ornl.gov/, in the “NW-BRaVE” collections in the Catalog of DataFed, ID 525611288 after registration.

- Project name: EcoKMER
- Project home page: https://github.com/NWBRaVE/EcoKMER
- Operating system(s): Platform independent
- Programming language: R (version 4.4.2 or later), Python (version 3.9.0 or later)
- License: Simplified BSD license

## Competing Interests

The authors declare that they have no competing interests.

## Funding

The research described in this paper is supported by the NW-BRaVE for Biopreparedness project funded by the U. S. Department of Energy (DOE), Office of Science, Office of Biological and Environmental Research, under FWP 81832. A portion of this research was performed on a project award (Enhancing biopreparedness through a model system to understand the molecular mechanisms that lead to pathogenesis and disease transmission) from the Environmental Molecular Sciences Laboratory, a DOE Office of Science User Facility sponsored by the Biological and Environmental Research program under Contract No. DE-AC05-76RL01830. Pacific Northwest National Laboratory is a multi-program national laboratory operated by Battelle for the DOE under Contract DE-AC05-76RL01830. This research used resources of the Oak Ridge Leadership Computing Facility at the Oak Ridge National Laboratory, which is supported by the Advanced Scientific Computing Research programs in the DOE Office of Science under Contract No. DE-AC05-00OR22725.

## Authors’ contributions

CGMJ, PW, MSC, DDP, conceived the research. CGMJ, PW, supervised the software development. AG, DL, CP, JB implemented the software. BN, JMD validated the software. OPL, NRB, RW, LA provided and curated the research data. MSC, DDP, acquired funding. OK provided access to computational resources. CGMJ, PW, AG, DL, CP wrote the initial draft. All authors participated in editing and reviewing the manuscript.

## Notes

https://github.com/NWBRaVE/EcoKMER

### Competing Interest Statement

The authors have declared no competing interest.

### Summary of Updates

Author name correction; Conner Philips -> Conner Phillips. Added a link to published OSTI page 'https://www.osti.gov/biblio/3412938' in the abstract.

